# Phytophthora palmivora establishes tissue-specific intracellular infection structures in the earliest divergent land plant lineage

**DOI:** 10.1101/188912

**Authors:** Philip Carella, Anna Gogleva, Marta Tomaselli, Carolin Alfs, Sebastian Schornack

## Abstract

The expansion of plants onto land was a formative event that brought forth profound changes to the Earth’s geochemistry and biota. Filamentous eukaryotic microbes developed the ability to colonize plant tissues early during the evolution of land plants, as demonstrated by intimate symbiosis-like associations in >400 million-year-old fossils. However, the degree to which filamentous microbes establish pathogenic interactions with early divergent land plants is unclear. Here, we demonstrate that the broad host-range oomycete pathogen *Phytophthora palmivora* colonizes liverworts, the earliest divergent land plant lineage. We show that *P. palmivora* establishes a complex tissue-specific interaction with *Marchantia polymorpha*, where it completes a full infection cycle within air chambers of the dorsal photosynthetic layer. Remarkably, *P. palmivora* invaginates *M. polymorpha* cells with haustoria-like structures that accumulate host cellular trafficking machinery and the membrane-syntaxin MpSYP13B but not the related MpSYP13A. Our results indicate that the intracellular accommodation of filamentous microbes is an ancient plant trait that is successfully exploited by pathogens like *P. palmivora*.

## INTRODUCTION

Plant-microbe associations are ubiquitous throughout the plant kingdom, which suggests that the ability to support microbial colonization occurred early during the evolution of land plants. Extensive evidence reveals that the first land plants were colonized by filamentous eukaryotic microbes, as several fossils from the Rhynie Chert (Early Devonian, 400-480 MYA) contain symbiosis-like microbial structures within ancient plant cells [1-4]. Indeed, symbiotic interactions with arbuscular mycorrhizal fungi are widespread among extant early land plants [5-8]. Moreover, recent studies suggest that early land plants and their algal predecessors were pre-adapted for symbiosis, as these organisms encode functionally equivalent homologs of core symbiosis signaling components [9,10]. In comparison, our understanding of how early divergent land plants interact with pathogenic microbes remains extremely limited.

Interactions between plants and filamentous eukaryotic microbes are often associated with the development of specialized microbial structures that protrude into plant cells. Such structures include finely-branched arbuscules of symbiotic arbuscular mycorrhizal fungi and digit, knob, or peg-like haustoria of oomycete and fungal pathogens [11,12]. Arbuscules and haustoria are both involved in the manipulation of host cell function to improve microbial colonization, however, symbiotic structures participate in the mutually beneficial exchange of resources while pathogenic structures function to suppress host immune responses and remove nutrients from the host [13,14]. Numerous host tissues and cells are capable of supporting filamentous microbes [13,15,16], which suggests that partially overlapping mechanisms are employed to accommodate symbiotic and pathogenic microbes. This has led to the idea that pathogens establish intracellular interfaces by exploiting host machinery designed to accommodate endosymbiotic structures. Whether this dynamic was established early during the co-evolution of plants and microbes is unknown. Extensive evidence has demonstrated that early land plants can accommodate arbuscules within their cells [7], whereas specialized pathogenic structures like haustoria have not been observed in these plants.

Bryophytes are a basal group of nonvascular, gametophyte-dominant (haploid) early land plants that include liverworts, hornworts, and mosses. Phylogenetic analyses often place liverworts basal to mosses and hornworts [17,18]. This suggests that liverworts represent the earliest divergent land plant lineage, although this has yet to yield a consensus among the field [19,20]. Many bryophytes are colonized by symbiotic microbes, however intracellular endosymbiotic structures (arbuscules) have only been observed in liverworts and hornworts [7,18]. Unfortunately, our understanding of plant-pathogen interactions in these plants is extremely limited. In comparison, several groups have described pathogenic interactions in the model moss *Physcomitrella patens* [21-23]. The colonization of moss tissues is associated with intracellular hyphal growth, however specialized infection structures similar to haustoria are not observed within moss cells [22]. An alternative bryophyte model system is therefore required to study intracellular interactions with pathogenic microbes. Liverworts are the most suitable candidate to accomplish this, given their ability to accommodate endosymbiotic structures and the recent establishment of molecular genetic tools in these plants [24].

In this study, we investigated plant-pathogen interactions in the model liverwort *Marchantia polymorpha*, which propagates clonally via small propagules (gemmae) or sexually when the sperm of male plants fertilize eggs housed in the archegonia of female plants. Both male and female plants have a thalloid body plan comprised of a ventrally located epidermis with rhizoids and scales, a central non-photosynthetic storage region, and a dorsal photosynthetic layer [25]. The photosynthetic layer of complex liverworts like *M. polymorpha* is made up of air chambers, which are walled enclosures that contain plastid-rich photosynthetic filaments and have open pores to facilitate gas exchange [25].

To determine if liverworts support intracellular colonization by a filamentous eukaryotic pathogen, we challenged *M. polymorpha* with *Phytophthora palmivora*, a broad host-range hemi-biotrophic oomycete pathogen with demonstrated virulence in root and leaf tissues of monocot and dicot hosts [26-28]. In general, *P. palmivora* infection begins when motile zoospores contact plant surfaces, causing the spores to encyst, germinate and form appressoria to penetrate the surface [29]. Upon accessing plant tissues, the biotrophic phase of *P. palmivora* colonization begins and the pathogen develops digit-like haustoria that protrude into plant cells for nutrient acquisition and the release of virulence effector proteins that manipulate the host [11]. Precise mechanisms for oomycete effector delivery/uptake remain to be clarified, however secreted effector proteins containing the RXLR motif translocate into host cells and interfere with immunity [30-32]. The pathogen eventually transitions to a necrotrophic phase, where it actively destroys plant tissues and completes its asexual lifestyle by releasing motile zoospores contained within sporangia [33].

Here, we demonstrate that *P. palmivora* preferentially colonizes the photosynthetic layer of liverworts and causes extensive disease. Molecular and microscopic analyses revealed that the biotrophic phase of *P. palmivora* colonization is associated with the upregulation of virulence effector molecules and the deployment of haustoria-like intracellular infection structures. Several endogenous host proteins accumulated at intracellular infection structures, including Rab GTPases and the homolog of a membrane-localized syntaxin associated with symbiosis in higher plants. Together, these markers clearly defined atypically branched infection structures and intracellular hyphae, revealing a prevalent intracellular phase of *P. palmivora* pathogenesis that is not commonly observed during interactions with higher plants. Furthermore, we demonstrate that *P. palmivora* requires liverwort air chambers to fully exploit *M. polymorpha*, as pathogen fitness is greatly reduced in air chamber-less *nop1* mutants.

## RESULTS

### Phytophthora palmivora colonizes the photosynthetic layer of Marchantia polymorpha

To determine whether liverworts support colonization by a broad host-range filamentous pathogen, we challenged 3-week-old *M. polymorpha* TAK1 (male) plants with zoospores of a tdTomato-expressing isolate of *P. palmivora* (accession P3914, derived from Arizona, U.S.A; ARI-td) and tracked pathogen growth and disease progression. Since *P. palmivora* colonizes multiple tissue types in higher plants [27,28,34], initial experiments were performed to determine whether colonization occurs in rhizoids or thalli of *Marchantia*. Rhizoid inoculations led to variable disease phenotypes, such that plants occasionally exhibited disease symptoms by 7 days post inoculation (dpi; Fig S1A). Confocal fluorescence microscopy indicated that rhizoids were mostly devoid of intracellular *P. palmivora* (ARI-td) growth, with rare instances of intracellular hyphae in damaged rhizoids (Fig S1B). In contrast, *Marchantia* thalli inoculated with ARI-td zoospores exhibited disease symptoms that increased in severity from 3-7 dpi (Fig 1A). Epifluorescence and confocal microscopy supported these observations, demonstrating the gradual spread of ARI-td hyphae across TAK1 thalli from 1-4 dpi (Fig 1B, Fig S2). Plants were completely colonized by 7 dpi, demonstrating that *M. polymorpha* thalli are highly susceptible to *P. palmivora*.

**Fig 1.**
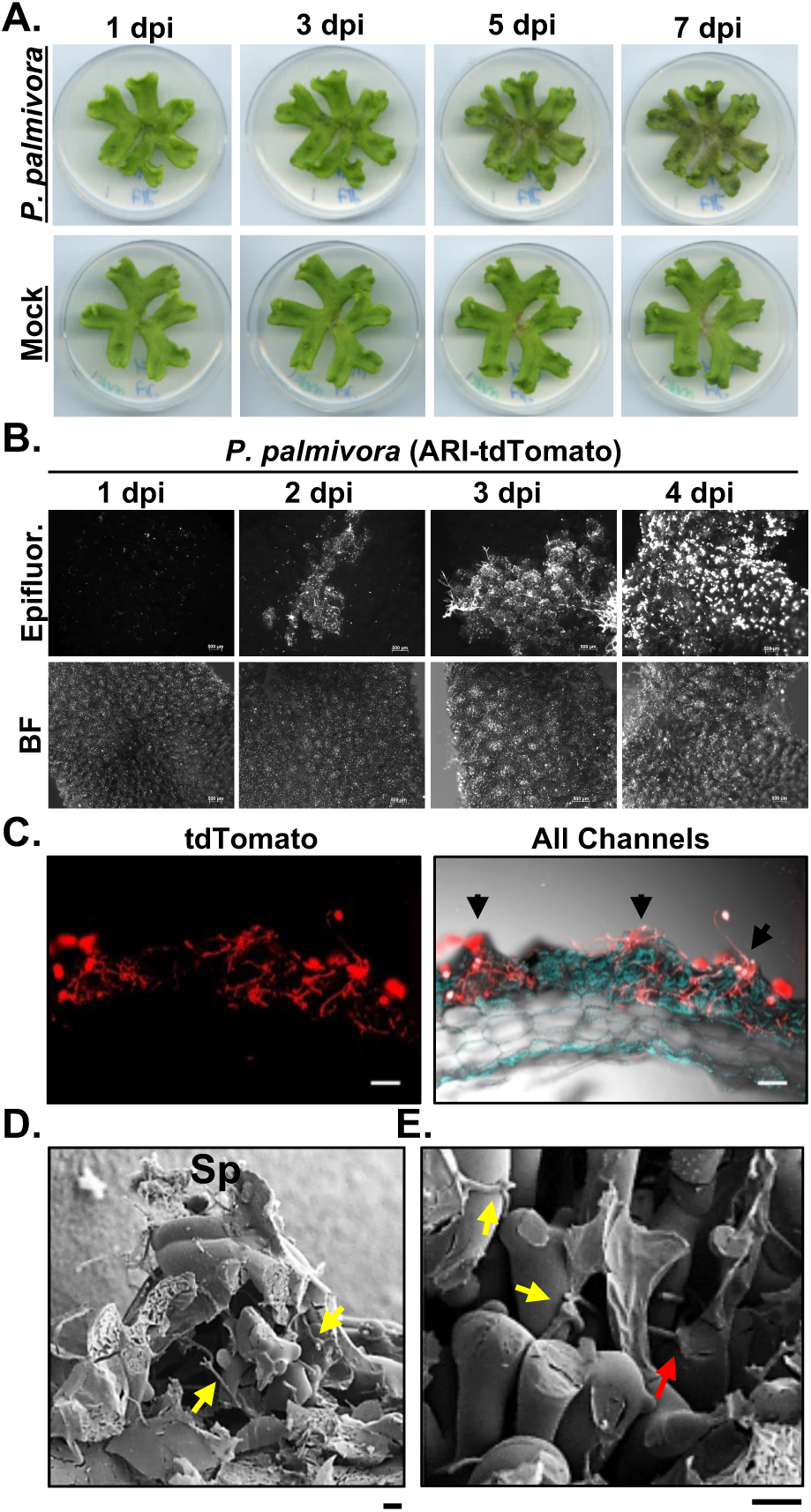
*Phytophthora palmivora* colonizes the photosynthetic layer of *Marchantia polymorpha*. **(A)** Disease symptoms of 3-week-old *M. polymorpha* TAK1 (male) thalli inoculated with *P. palmivora* ARI-td zoospores or water (mock) over a 7-day time course (dpi - days post inoculation). **(B)** Epifluorescence microscopy demonstrating the spread of *P. palmivora* growth across TAK1 thalli from 1-4 dpi. Epifluorescence (Epifluor.) from the pathogen is displayed alongside bright field (BF) images. Scale bars = 500 μm. **(C)** Confocal fluorescence microscopy of sectioned TAK1 thalli infected with *P. palmivora* at 7 dpi. Z-stack projections of red fluorescence from the pathogen is displayed alone (tdTomato) or merged with all channels (bright-field and plastid auto-fluorescence in turquoise). Arrows indicate air pores. Scale bars = 100 μm. **(D-E)** CryoSEM (scanning electron microscopy) of TAK1 thalli colonized by *P. palmivora* at 7 dpi. (D) Mechanically fractured air chamber demonstrating hyphal growth within the chamber (yellow arrows) and sporangia (Sp) at the air pore. (E) Intercellular (yellow arrows) and intracellular (red arrow) associations between *P. palmivora* hyphae and photosynthetic filaments within *M. polymorpha* air chambers. Scale bars = 20 μm. Experiments (A-E) were performed at least three times with similar results.

Confocal fluorescence microscopy demonstrated that *P. palmivora* colonized the surface of *M. polymorpha* thalli in a discrete manner that overlapped with air chamber morphology (Fig S2). We therefore analyzed cross sections of infected *M. polymorpha* thalli, which revealed high levels of colonization within air chambers at 7 dpi (Fig 1C). Pathogen growth was largely limited to the photosynthetic layer, although intercellular hyphae were observed in the non-photosynthetic storage tissue of plants with ongoing necrosis (Fig S3). In support of these findings, experiments performed using additional liverwort species (*Lunularia cruciata* and *M. paleacea*) similarly exhibited colonization within air chambers (Fig S4A). Cryo-SEM (scanning electron microscopy) of colonized *Marchantia* thalli demonstrated *P. palmivora* sporangia and hyphae traversing through the central pores of air chambers (Fig 1D) and hyphal growth was observed within air chambers (Fig 1D and 1E). *P. palmivora* hyphae often associated with photosynthetic filament cells and were sometimes observed to penetrate them (Fig 1E). Together, our results indicate that *P. palmivora* preferentially colonizes the air chambers of *M. polymorpha*.

Natural diversity of pathogen isolates has classically been used to probe host-microbe interactions to identify loci or mechanisms important for plant resistance to disease. To explore this paradigm in *Marchantia*, we assessed the ability of several *P. palmivora* isolates to colonize TAK1 thalli (Fig S5). Disease progression was similar to ARI-td for the majority of strains tested, with the exception of the MAZI (P6375, Malaysia) and TAZI (P6802, Thailand) isolates that displayed an accelerated disease progression (Fig S5). We also tested compatibility between *Marchantia* and *P. infestans*, the causal agent of potato and tomato late blight. TAK1 thalli infected with *P. infestans* (Pi-88069-td) were asymptomatic over a 7-day infection time course, similar to mock-treated plants (Fig S6). In contrast, TAK1 thalli inoculated with *P. palmivora* (ARI-td) zoospores were highly susceptible to pathogen ingress. These results demonstrate that *P. infestans* is unable to overcome preexisting or induced barriers to colonization in *Marchantia*, unlike several *P. palmivora* isolates that cause extensive disease.

### P. palmivora establishes a biotrophic interaction with host-intracellular oomycete hyphae

The colonization of higher plant tissues by *P. palmivora* is associated with distinct transitions in pathogen lifestyle. These transitions are typically characterized by the presence of specialized penetration/infection structures and clear shifts in the *P. palmivora* transcriptome [28]. We first performed confocal fluorescence microscopy to document the pathogen lifestyle transitions that occur during the colonization of TAK1 thalli (Fig 2A). At 1 dpi, hyphae of germinated zoospores appeared to wander along the surface of the thallus without developing appressorial penetration structures (Fig 1B, Fig 2A, Fig S2). The colonization of plant tissues occurred by 2 dpi and was associated with the development of biotrophic intracellular infection structures. We observed typical digit-like haustoria, however, highly branched intracellular structures and intracellular hyphae were observed in greater abundance (Fig 2A). For every digit-like haustorium, there were approximately four times as many branched intracellular structures and twice as many instances of intracellular hyphal growth. Intracellular structures were also observed during interactions with *L. cruciata* and *M. paleacea*, however digit or knob-like haustoria appeared to be more prevalent than branched infection structures (Fig S4B). Sporangia were observed by 3-4 dpi (Fig 2A), indicating that *P. palmivora* completes a full infection cycle in *Marchantia*.

**Fig 2.**
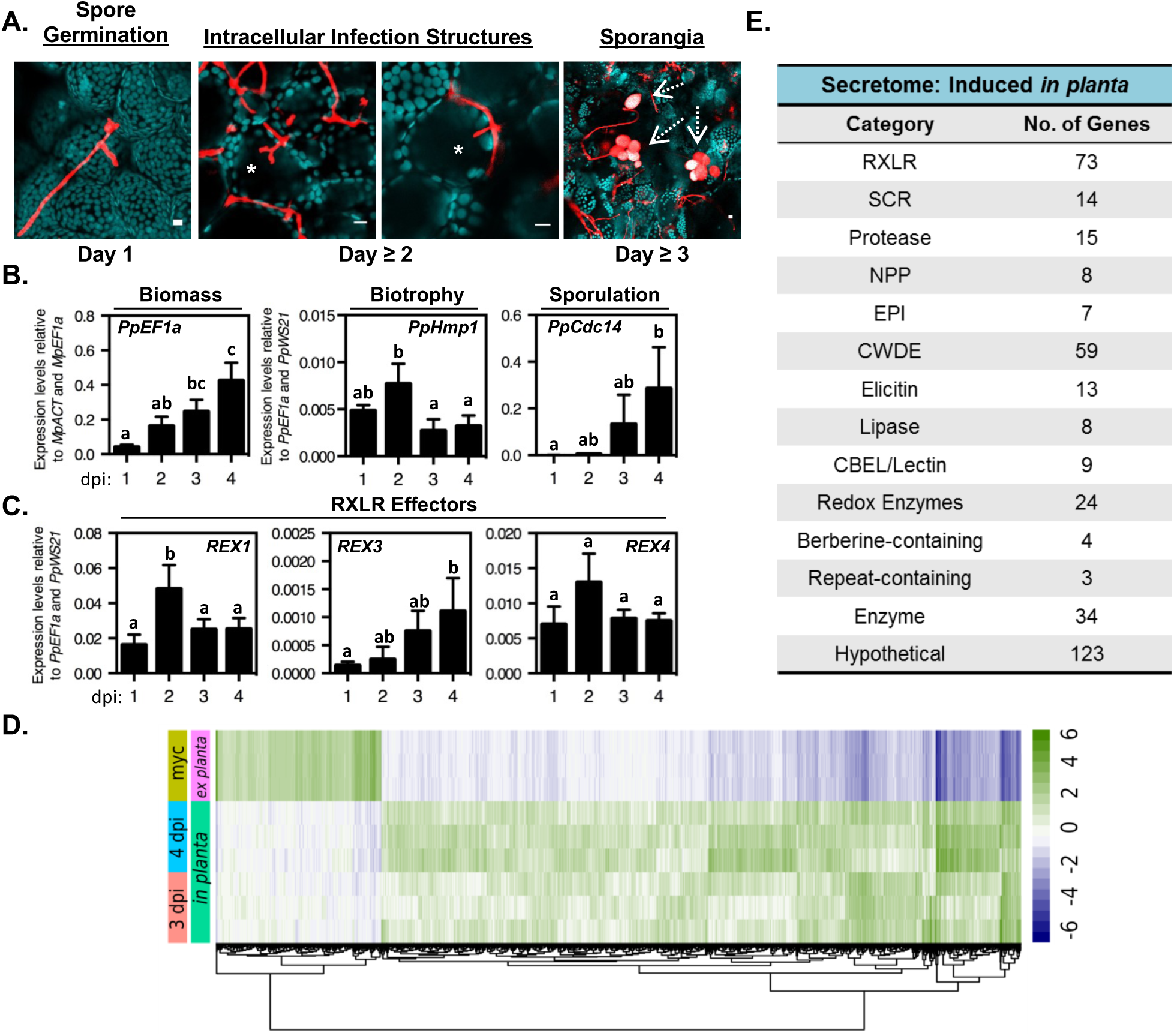
*Phytophthora palmivora* establishes a biotrophic interaction and completes a full infection cycle in *Marchantia*. **(A)** Confocal fluorescence microscopy demonstrating key morphological transitions in *P. palmivora* lifestyle during the colonization of TAK1 plants from 1-3 days post inoculation (dpi). Z-stack projections of pathogen fluorescence merged with plastid auto-fluorescence (turquoise). Intracellular infection structures are denoted by an asterisk (*). Sporangia are indicated by dashed arrows. Scale bars = 10 μm. **(B)** Quantification of *P. palmivora* lifestyle marker genes during the colonization of TAK1 thalli from 1-4 dpi via qRT-PCR analysis. Pathogen biomass (*PpEFIa*) was quantified relative to *M. polymorpha* biomass markers (*MpACT* and *MpEFIa*). Biotrophy (*PpHmpI*) and sporulation (*PpCdc14*) marker genes were quantified relative to pathogen biomass controls (*PpEFIa* and *PpWS21*). Different letters signify statistically significant differences in transcript abundance (ANOVA, Tukey’s HSD p < 0.05). **(C)** Quantification of *P. palmivora* RXLR effector gene transcripts during the colonization of TAK1 thalli from 1-4 dpi via qRT-PCR analysis. RXLR effector (*REX1, REX3, REX4*) gene expression was quantified relative to *P. palmivora* biomass (*PpEFIa* and *PpWS21*). Different letters signify statistically significant differences in transcript abundance (ANOVA, Tukey’s HSD p <0.05). **(D)** *P. palmivora* transcriptome. Hierarchical clustering of differentially expressed genes between *in planta* (3 and 4 dpi) and axenically grown mycelium transcriptomes (LFC ≥ 2, p.value ≤ 10^−3^). Rlog-transformed counts, median-centered by gene are shown. **(E)** *P. palmivora* secretome. Summary of functional categories of 394 genes encoding putative secreted proteins up-regulated during *P. palmivora* infection of *M. polymorpha*. Experiments (A-C) were performed at least three times with similar results.

To further clarify the timing of pathogen lifestyle transitions during the *Marchantia-Phytophthora* interaction, we monitored the expression of lifestyle-associated *P. palmivora* marker genes over a four-day infection time course by qRT-PCR analysis (Fig 2B). Levels of the *P. palmivora* biomass marker *PpEF1a* significantly increased over time, which is consistent with our microscopic analyses (Fig 1B, Fig S2). The biotrophy-specific marker gene *PpHmp1* (*Haustoria Membrane Protein1* [35]) peaked in expression at 2 dpi, while the sporulation-specific cell cycle marker gene *PpCdc14* [36] was induced by 3-4 dpi. These results are consistent with our microscopy data that demonstrates intracellular infection structures at 2 dpi and sporangia at 3-4 dpi (Fig 2A). Collectively, our results indicate that the *P. palmivora* infection cycle in *Marchantia* begins with spore germination and hyphal growth at 1 dpi, followed by a biotrophic phase that includes intracellular infection structures at 2-3 dpi, and ends with the completion of the pathogen’s asexual life-cycle after 3 dpi.

The transcriptional induction of pathogen genes encoding secreted effector proteins is a hallmark of *Phytophthora*-angiosperm interactions. Among these secreted proteins, the RXLR class of effectors are believed to act within host cells to suppress immunity and enhance pathogen growth [30-32]. To determine if *P. palmivora* upregulates RXLR effectors during the colonization of *Marchantia*, we analyzed the expression profiles of the *P. palmivora* RXLR effectors *REX1, REX3*, and *REX4* [28]. Significant upregulation of *REX1* transcripts occurred during the biotrophic phase at 2 dpi and *REX3* levels increased throughout infection to a maximum observed at 4 dpi (Fig 2C). *REX4* expression peaked at 2 dpi, although this was not statistically significant in all experiments. *REX* expression profiles were similar to those observed in colonized *N. benthamiana* roots [28], demonstrating that *P. palmivora* upregulates RXLR effector transcripts during the colonization of *Marchantia* thalli.

In order to assess broad transcriptional changes in *P. palmivora* during liverwort colonisation, we performed RNA-sequencing of infected thalli (3 and 4 dpi) and axenically grown *P. palmivora* mycelia, which served as an *ex planta* control. Differential expression analysis revealed 3601 up-regulated and 932 down-regulated *P. palmivora* genes expressed *in planta* relative to the *ex-planta* control (absolute LFC ≥ 2, adjusted *p-value* < 10^−3^) (Fig 2D; Table S1). qRT-PCR analysis validated a subset of colonization-induced *P. palmivora* genes, which were either upregulated throughout colonization or were induced late during infection (Fig S7). Amongst all up-regulated genes, 394 (11%) were predicted to encode putative secreted proteins with RXLR effectors and cell wall degrading enzymes being the most abundant functional categories (Fig 2E). Taken together these results suggest that *P. palmivora* infection of *M. polymorpha* involves large-scale transcriptional induction of genes typical of a compatible interaction, including effectors and hydrolytic enzymes.

### Host Cellular Responses to Intracellular Infection Structures

The accommodation of intracellular hyphal structures requires reorganization of the host cell and the biogenesis of novel membranes to separate the host cell from the pathogen. Moreover, plants responding to invading microbes often accumulate callose, a β-1,3-glucan associated with cell wall strengthening. We therefore applied callose staining and live-cell imaging of fluorescently tagged proteins labelling membrane compartments to investigate the subcellular changes associated with accommodating intracellular *P. palmivora* structures in *Marchantia*. An extended callosic envelope is characteristic of dysfunctional intracellular structures while functional interfaces display callosic collars that are limited to the neck region or no callosic depositions at all [37,38]. Aniline blue staining of ARI-td-infected TAK1 thalli revealed that callose deposition was limited to the peripherally-located neck region of intracellular infection structures (Fig 3A, Fig S8), suggestive of functional *P. palmivora-Marchantia* interfaces. Next, we localized homologs of host Rab GTPases that label oomycete haustoria in higher plants and are believed to direct vesicle delivery to host-pathogen interfaces [39]. We generated transgenic plants that constitutively overexpress mCitrine-MpRabG3c or mCitrine-MpRabA1e, since these proteins strongly localize to *P. infestans* haustoria in *N. benthamiana* leaves [39,40]. In the absence of *P. palmivora* colonization, mCitrine-MpRabG3c and mCitrine-MpRabA1e localized to the tonoplast and to endosomes, respectively (Fig S9), which is consistent with RabG3c and RabA1e localization in higher plants [40,41]. During infection, we observed strong labelling of intracellular infection structures and intracellular hyphae by mCitrine-MpRabG3c (Fig 3B) as well as mCitrine-MpRabA1e (Fig 3C) at 3 dpi. Together, our results reveal a conserved host cellular response to invasive haustoria-like intracellular infection structures and hyphae during the colonization of *Marchantia* cells by *P. palmivora*.

**Fig 3.**
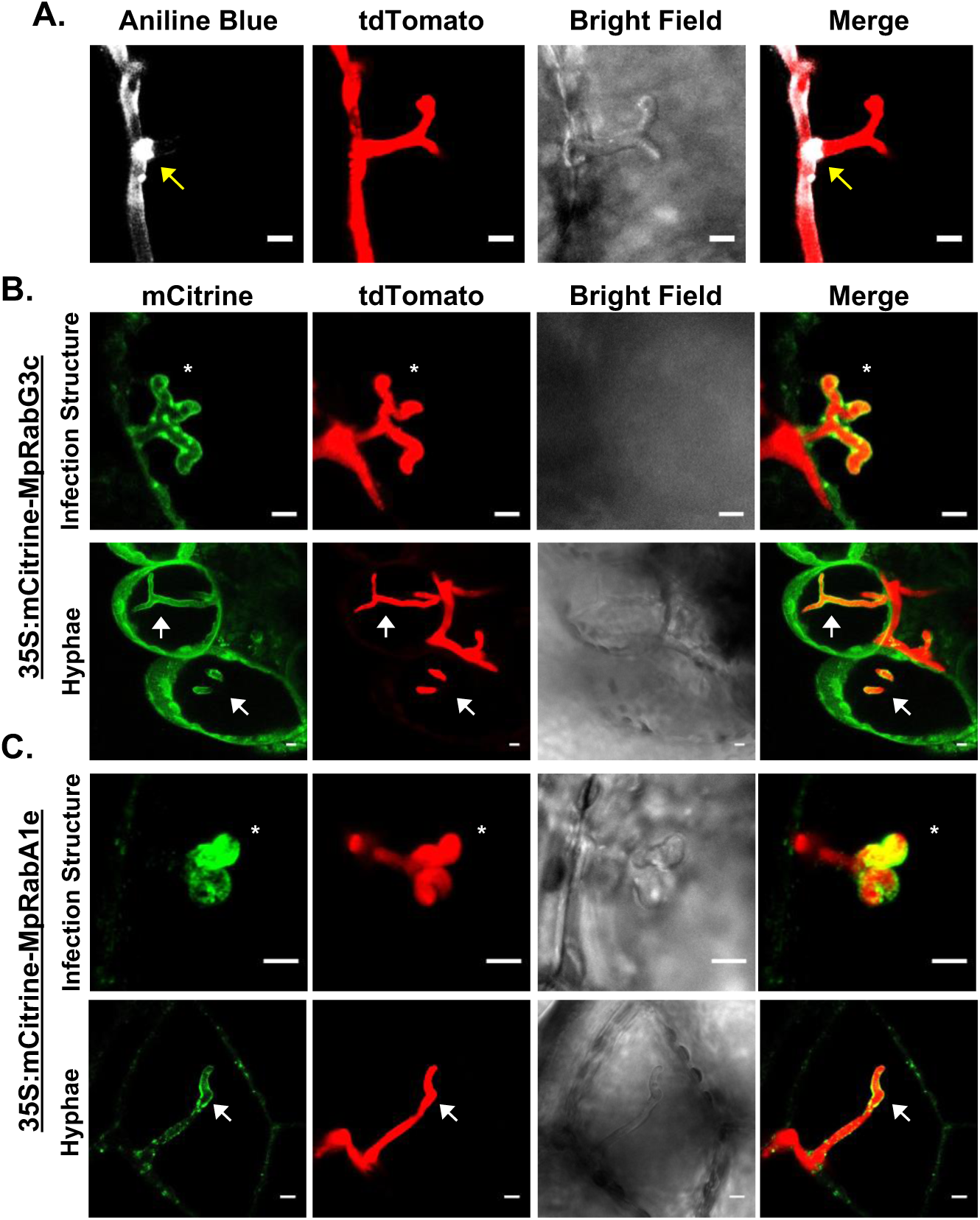
Host Cellular Responses to Invading Oomycete Structures. **(A)** Detection of callose deposition at *P. palmivora* ARI-td intracellular infection structures at 3 days post inoculation (dpi). Arrows indicate callose deposition at the peripheral neck region of the invading intracellular infection structure (denoted by an asterisk *). Scale bar = 5 μm. **(B)** MpRabG3c co-localization with invading *P. palmivora* ARI-td structures in 35S:mCitrine-MpRabG3c plants at 3 dpi. Infection structures are indicated with asterisks (*) while intracellular hyphae are denoted by arrows. Z-stack projections are displayed. Scale bars = 5 μm. **(C)** MpRabA1e co-localization with invading *P. palmivora* ARI-td structures in 35S:mCitrine-MpRabA1e plants at 3 dpi. Z-stack projections are displayed. Scale bars = 5 μm. Experiments (A-C) were performed three times with similar results.

### A colonization-induced host syntaxin is targeted to intracellular infection structures

Plasma membrane-resident syntaxins mediate exocytosis and have been associated with penetration structures of symbiotic and pathogenic microbes in several higher plant systems [42,43]. To determine if symbiosis-associated membrane syntaxins function at intracellular infection structures, we monitored the expression levels of *Marchantia SYP13* family homologs during infection and assessed MpSYP13 localization at pathogen interfaces using mCitrine-tagged reporter lines [44]. Expression levels of *MpSYP13A* were not affected during colonization with *P. palmivora*, while *MpSYP13B* displayed significant upregulation from 2-4 dpi (Fig 4A). In addition, mCitrine-MpSYP13B strongly labelled intracellular infection structures while maintaining localization at the cell periphery. Conversely, mCitrine-MpSYP13A localization was unaffected by the presence of intracellular infection structures (Fig 4B). We observed a number of distinct MpSYP13B localization patterns, including focal accumulation proximal to hyphal buds of early developing intracellular infection structures, complete labelling around intracellular hyphae, and localization to membrane domains of branched intracellular infection structures (Fig 4C). These results demonstrate that the colonization of *Marchantia* by *P. palmivora* includes a complex intracellular phase that recruits the MpSYP13B syntaxin to specific extrainvasive hyphal domains.

**Fig 4.**
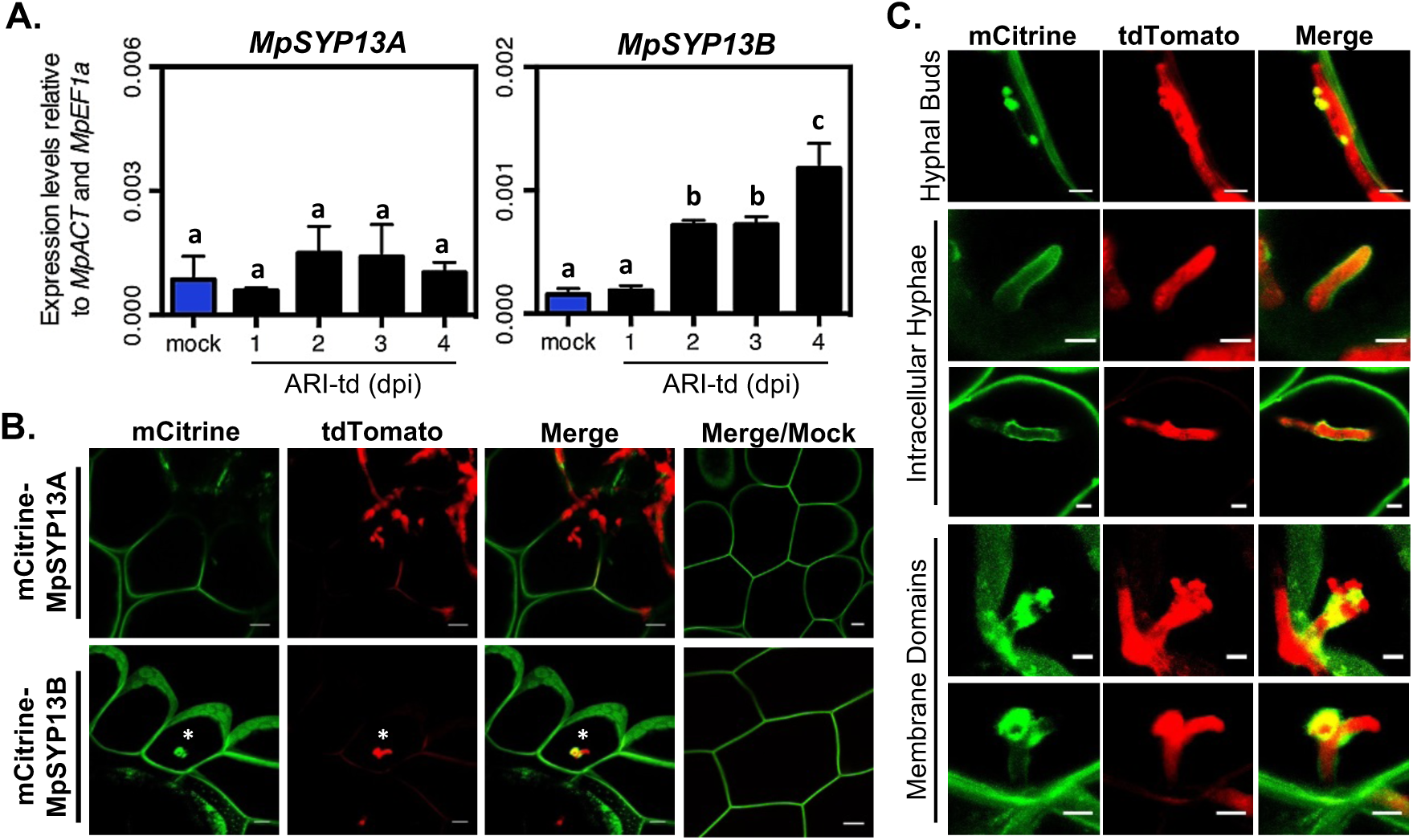
A colonization-induced host syntaxin accumulates at intracellular infection structures. **(A)** qRT-PCR analysis of *MpSYP13A* and *MpSYP13B* transcripts in mock-treated or *P. palmivora*-colonized (ARI-td) TAK1 plants from 1-4 days post inoculation (dpi). Expression values are shown relative to internal *MpACT* and *MpEF1a* controls. Different letters signify statistically significant differences in transcript abundance (ANOVA, Tukey’s HSD p < 0.05). **(B)** Confocal fluorescence microscopy demonstrating mCitrine-MpSYP13A/B localization in cells containing *P. palmivora* (ARI-td) intracellular infection structures at 3 dpi. Asterisks denote intracellular infection structures. Scale bars = 10 μm. **(C)** Patterns of mCitrine-MpSYP13B localization in *P. palmivora*-colonized (ARI-td) plants, including a close-up image of the structure displayed in (B). Scale bars = 5 μm. Experiments (A-C) were performed three times with similar results.

### Co-option of Marchantia air chambers for pathogen colonization

The colonization dynamics described thus far have occurred in epidermal cells and within air chambers of the photosynthetic layer of *M. polymorpha*. In *Marchantia*, loss of function mutations in the E3 ubiquitin ligase NOPPERABO (NOP1) result in the development of a photosynthetic layer that lacks air chambers entirely [45]. To determine if air chambers facilitate *P. palmivora* colonization, we compared the colonization phenotypes of wild-type TAK1 and air chamber-less *nop1* mutants (Fig 5). As expected, *P. palmivora* ARI-td zoospores inoculated onto TAK1 thalli caused extensive disease symptoms by 7 dpi and continued to 14 dpi when plants were essentially dead. In comparison, air chamber-less *nop1* mutants displayed reduced disease symptoms, with plants remaining relatively healthy throughout the experiment (Fig 5A). To support these observations, we quantified the expression of pathogen-specific marker genes indicative of biomass (*PpEF1a*) and sporulation (*PpCdc14*) by qRT-PCR. Compared to wild-type TAK1, pathogen biomass and sporulation were both significantly reduced in *nop1* mutants (Fig 5B), which indicates that *P. palmivora* fitness is largely dependent on the presence of air chambers. This was further supported by microscopic analyses of *P. palmivora* growth in TAK1 compared to *nop1* plants. Confocal fluorescence microscopy of sectioned thalli revealed extensive hyphal growth within TAK1 air chambers at 7 dpi, whereas *nop1* plants displayed only a thin layer of *P. palmivora* hyphae on the dorsal epidermis (Fig 5C). Moreover, cryo-SEM micrographs confirmed that hyphae travel from chamber-to-chamber during the colonization of TAK1 thalli, with widespread hyphal growth observed within air chambers (Fig 5D). In comparison, a network of surface hyphae was observed on *nop1* thalli and collapsed epidermal cells containing *P. palmivora* hyphae were occasionally detected late during infection (Fig 5E). In support of this, confocal fluorescence microscopy identified invasive ARI-td hyphal growth within *nop1* epidermal cells by 3 dpi (Fig 5F). Taken together, our results demonstrate that *nop1*-dependent changes in thallus architecture impair *P. palmivora* proliferation, suggesting a key role for air chambers in supporting pathogen colonization.

**Fig 5.**
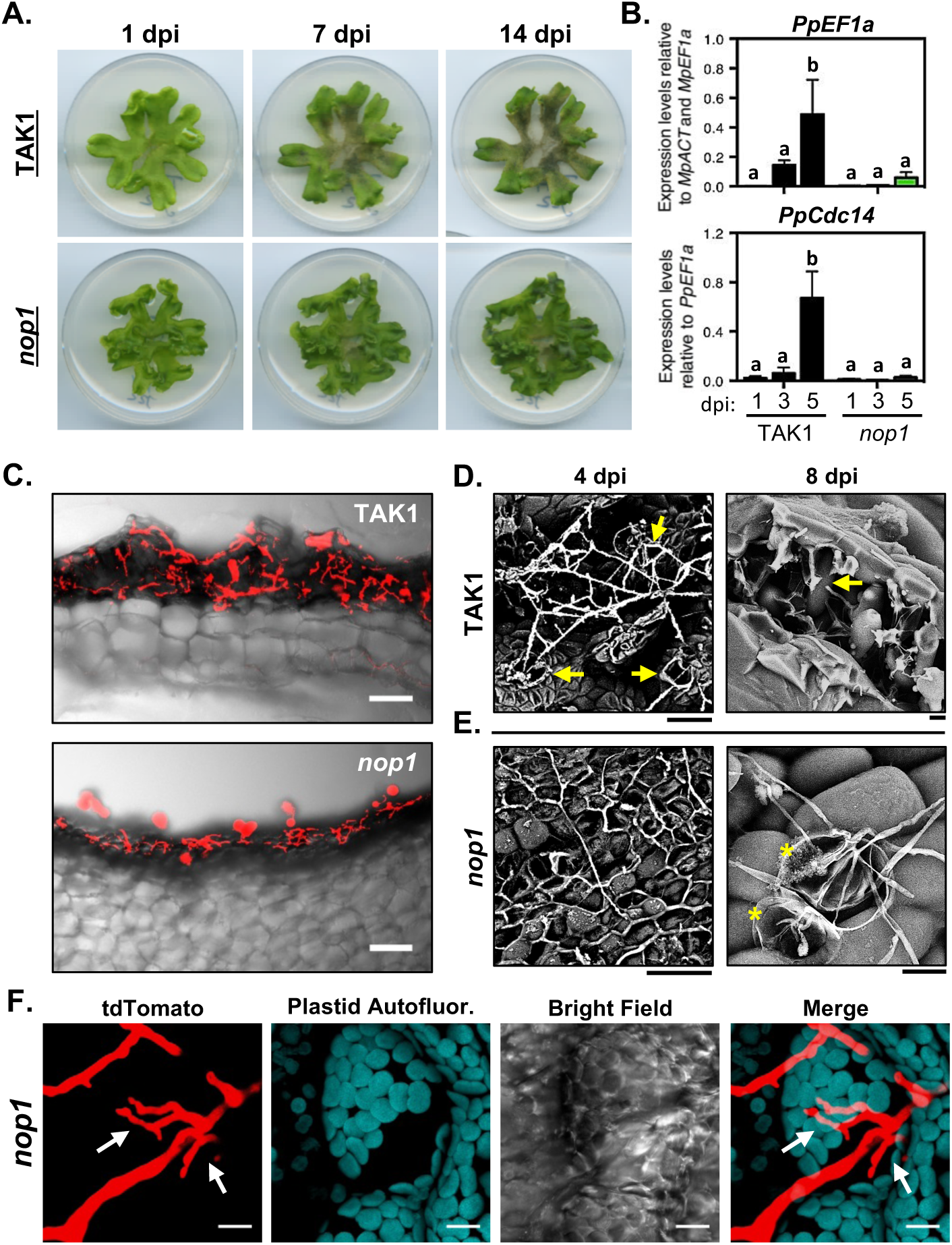
*Phytophthora palmivora* requires air chambers for the successful colonization of *Marchantia polymorpha* thalli. **(A)** Disease symptoms of 3-week-old *M. polymoprha* TAK1 (wild-type) and *nop1* mutant plants inoculated with *P. palmivora* ARI-td zoospores at 1, 7, and 14 days post inoculation (dpi). **(B)** Quantification of *P. palmivora* biomass (*PpEF1a*) and sporulation (*PpCdc14*) marker genes during the colonization of wild-type TAK1 and *nop1* plants at 1, 3, and 5 dpi. *PpEF1a* expression was quantified relative to *M. polymorpha* biomass markers (*MpACT* and *MpEF1a*). *PpCdc14* was quantified relative to pathogen biomass (*PpEF1a*). Different letters signify statistically significant differences in transcript abundances (ANOVA, Tukey’s HSD p< 0.05). **(C)** Confocal fluorescence microscopy of sectioned TAK1 and *nop1* thalli infected with *P. palmivora* ARI-td at 7 dpi. Micrographs display merged z-stack projections of red pathogen fluorescence and bright field images. Scale bars = 100 μm. **(D-E)** Cryo-SEM (scanning electron microscopy) of TAK1 and *nop1* thalli infected with *P. palmivora* at 4 or 8 dpi. (D) Micrographs of TAK1 plants demonstrate ARI-td hyphae travelling between air pores (yellow arrows) at 4 dpi and hyphal growth (yellow arrow) within air chambers at 8 dpi. (E) Micrographs of *nop1* plants demonstrate a network of surface hyphae at 4 dpi and collapsed epidermal cells (asterisks) are sometimes observed at 8 dpi. Scale bars = 20 μm. **(F)** Confocal fluorescence microscopy demonstrating invasive ARI-td hyphal growth in *nop1* epidermal cells at 3 dpi. Micrographs display z-stack projections of red pathogen fluorescence (tdTomato), plastid autofluorescence, bright field images, and plastid and tdTomato fluorescence merged together. Scale bars = 10 μm. Experiments were performed three times (A,B,C,F) or twice (D,E) with similar results.

## DISCUSSION

In this study, we demonstrate that the broad host-range oomycete *P. palmivora* colonizes the photosynthetic layer of non-vascular liverworts and forms intracellular hyphal structures resembling a biotrophic interaction. This was supported by pathogen effector gene expression and by the presence of intracellular hyphal structures that display a biotrophy-associated callose distribution pattern and associate with host proteins typical of biotrophic interactions. Considered alongside observations of arbuscules in liverworts and hornworts [7,18,46], the data indicate that descendants of the earliest diverging land plants have the capacity to accommodate the biotrophic structures of filamentous eukaryotes within their cells.

The colonization of *Marchantia* thalli by *P. palmivora* was largely limited to the photosynthetic layer, with prolific hyphal growth within and between air chambers. Colonization assays with the air chamber-less *nop1* mutant revealed that these structures largely contribute to disease susceptibility in *Marchantia*. This is likely due to the fact that air chambers offer photosynthetic filament cells rich in carbohydrates and provide a microenvironment with stable humidity, which is ideal for water molds such as *Phytophthora*. Our results underscore the need for plants to protect intercellular spaces important for gas exchange. Such adaptations are present in higher plants, where analogous intercellular spaces (spongy mesophyll) are protected by stomata. In many cases, the early detection of pathogens causes stomatal closure to prevent the colonization of the intercellular space [47]. However, several pathogens have developed mechanisms to circumvent this strategy, either by accessing the tissue by other means or by the direct regulation of stomatal guard cells [48]. Most filamentous pathogens bypass stomata by penetrating host surfaces using appressoria, a penetration structure that does not appear to play a major role during the *P. palmivora-Marchantia* interaction. Rather, air chambers were accessed via intra-or intercellular hyphal growth, or through constitutively open pores. *P. palmivora* sporangia often emanated from air pores, similar to observations of sporangia traversing stomata in angiosperms. The sporophytes of mosses and hornworts contain stomata, but liverworts do not. How these early forms of stomata function in response to biotic cues is unknown, however elevated levels of CO2 do not alter bryophyte stomata as they do in angiosperms [49]. It is therefore possible that interactions with pathogenic microbes contributed to the development of stomata in early land plants. Future comparative studies are required to explore this relationship between microbial colonization and the development and regulation of pores involved in gas exchange.

The cell walls and surfaces of extant early land plants are less complex than those of higher plants [50]. How this impacts interactions with pathogenic or symbiotic microbes remains to be explored. It is tempting to speculate that simple cell walls are susceptible to invading microbes, however this may not be the case. For example, previous work in *Marchantia* demonstrated a differential capacity for powdery mildew spore establishment, such that *Erysiphe trifoliorum* spores germinated but were destroyed on *M. polymorpha* surfaces while *Oidium neolycopersici* survived and developed appressoria [51]. Whether *O. neolycopersici* penetrates into *M. polymorpha* cells and establishes infection was not explored by the authors. A recent study showed that *P. infestans* and *P. capsici* hyphae penetrate protonemal cells during non-host interactions in moss, whereas colonization by *P. palmivora* and *P. sojae* was rarely detected [22]. Both *P. infestans* and *P. capsici* fail to establish biotrophy in moss, as haustoria-like structures were never observed [22]. In contrast, we demonstrated the colonization of *Marchantia* thalli by *P. palmivora* rather than *P. infestans*, which was associated with the development of haustoria-like infection structures. Susceptibility to *P. palmivora* was greatly enhanced when thalli rather than rhizoids were inoculated with zoospores. In addition, intercellular hyphal growth within the central storage parenchyma was less prevalent than growth within air chambers, although plants eventually succumb to disease and are likely colonized throughout. Based on these observations, we hypothesize that the central storage region is resistant to pathogen colonization and requires prolific biotrophic growth within air chambers and a subsequent switch to necrotrophy for colonization. In contrast, the colonization of liverworts by symbiotic fungi is abundant within the central storage region but does not extend to the photosynthetic layer [5]. Collectively, these observations suggest that characteristics of early land plant tissues and surfaces act as barriers to microbial growth. Future efforts to identify the structural, genetic, or chemical components responsible for these phenotypes may reveal interesting similarities and differences to higher plants.

We identified several aspects of *P. palmivora* colonization that differ in *Marchantia* compared to interactions with angiosperms. Appressorial penetration structures were not prevalent in *Marchantia* as they are in higher plants [27,28]. The development of oomycete and fungal appressoria require cues such as hydrophobicity and the perception of cutin monomers characteristic of host cuticles [52,53]. The lack of *P. palmivora* appressoria on *Marchantia* surfaces suggests that key differences in cuticle composition impact pathogen development on liverwort surfaces. Indeed, bryophytes are poikilohydric plants that contain simple cuticle-like layers which afford desiccation tolerance and allows for water absorption through plant bodies [54]. Interestingly, the fungus *O. neolycopersici* develops appressoria on *Marchantia* [51], which may suggest differences in the detection of host surfaces/chemicals compared to *P. palmivora*. The biotrophic phase of the *P. palmivora-Marchantia* interaction was associated with highly branched intracellular infection structures and intracellular hyphae, whereas interactions in higher plants are associated with digit-like haustoria and intercellular hyphal growth. Several genetic markers for biotrophy were expressed during interactions with *Marchantia*, which suggests the presence of a biotrophic stage despite the under-representation of typical digit-haustoria. Branched intracellular infection structures were labelled by host proteins that are typically associated with the extra-haustorial matrix/membrane [39,40], which suggests a function in biotrophy. Interestingly, a range of intracellular infection structure morphologies were observed in different liverworts, with *L. cruciata* and *M. paleacea* also demonstrating the presence of digit and knob-like haustoria, respectively. This suggests that characteristics of liverwort cells influence the development of intracellular infection structures. Whether this applies to other bryophyte species remains to be determined, as intracellular infection structures have not been described in mosses and hornworts.

The membrane-localized SYP132 syntaxin family is associated with symbiosis in higher plants. In *Medicago truncatula*, the presence of symbiotic microbes induces alternative splicing of the *SYP132* locus to produce SYP132a, which in turn labels the infection threads and symbiosomes of nitrogen-fixing rhizobacteria [55,56]. Moreover, silencing of *SYP132a* suppressed mycorrhizal colonization in Medicago roots, demonstrating the importance of SYP132a in promoting multiple symbiotic associations [56]. We similarly observed colonization-induced upregulation and interface localization of a specific membrane syntaxin variant in *Marchantia* (MpSYP13B), however this was afforded by homologous gene copies rather than alternative splicing. These data suggest that the evolution of membrane-localized plant syntaxins was influenced by interactions with symbiotic and pathogenic microbes, perhaps diverging for roles in general secretory processes or for the accommodation of intracellular microbial structures. We hypothesize that MpSYP13B represents a specialized syntaxin that evolved to label intracellular structures in liverworts (this work) similar to SYP132a function in Medicago [56].

Early divergent descendants of the first land plants can accommodate biotrophic filamentous microbes as diverse as oomycetes (this study) and mycorrhizal fungi [5]. Our data demonstrate that liverworts encode proteins that respond and localize to intracellular infection structures and pathogen hyphae, which suggests that the molecular toolkit required to support intracellular colonisation by eukaryotic microbes emerged early during land plant evolution. Previous work has established conserved genetic components required for endosymbiosis, with experimental evidence demonstrating the presence of functionally conserved homologs of key symbiosis genes in bryophytes and algae [9,10]. Here we complement this work by demonstrating the intracellular colonization of liverworts by a filamentous pathogen that likely hijacks this machinery to cause disease. Together, these studies suggest an ancient origin for the evolution of microbial accommodation in plants, which is supported by observations of highly branched microbial structures and hyphae inside fossilized plant cells [1-4]. In liverworts, we observed highly branched haustoria-like structures and prolific intracellular hyphal growth that could appear symbiosis-like in morphology. We therefore suggest caution in attributing the presence of branched intracellular structures to symbiotic interactions based solely on the interpretation of fossilized microbial structures or microscopy-based analyses. Our work further underscores the need to consider pathogenic microbes alongside symbiotic counterparts when discussing the evolution of microbial accommodation processes in land plants.

## MATERIALS AND METHODS

### Plant Growth

The plants used in this study are described in Table S2. All plants were cultivated from gemmae under axenic conditions. *M. polymorpha* and *M. paleacea* were grown on ½ strength MS (Murashige and Skoog) media (pH 6.7) supplemented with B5 vitamins under continuous light (70 μE m^−2^ s^−1^) at 22 °C. *L. cruciata* were grown in short-day photoperiod conditions (9 hrs light) at 20 °C on M media [10].

### Pathogen Growth and Infection Assays

Pathogen strains used in this study are described in Table S3. *P. palmivora* ARI-td mycelia were maintained by routine passaging on RSA (Rye Sucrose Agar) plates and zoospores were collected from 7-10 day old cultures on V8 plates as previously described [27]. *P. infestans* zoospores were collected similar to *P. palmivora*, except the pathogen was grown solely on RSA plates incubated at 18 °C. Colonization experiments were performed by applying 10 uL droplets of a zoospore suspension inoculum (10^5^ zoospores mL^−1^) along the thallus or directly onto rhizoids of 3 week-old *M. polymorpha* and *M. paleacea* plants. Slower growing *L. cruciata* liverworts grown from gemmae were infected 6-8 weeks post plating.

### Microscopy

Epifluorescence microscopy was conducted using a Zeiss Axioimager stereo-fluorescence microscope with a DsRED filter. All images were processed with AxioVision software (Rel 4.8). Confocal fluorescence microscopy was performed using a Leica TCS SP8 equipped with HyD detectors. Settings for pathogen detection (tdTomato) are described in [27], mCitrine in [44], and aniline blue in [57]. All experiments were performed at least 3 times with similar results. Images were collected from at least 3 independent plants in at least 2 separate infection sites per plant. Cross sections of a 200-300 μm thickness were prepared from agarose (3%) embedded samples using a vibratome. All images were processed using ImageJ. Cryo-scanning electron microscopy (SEM) was performed essentially as described in [58], except that ARI-td-infected *M. polymorpha* tissue was used. All samples were sublimated for 30-60 seconds. Where indicated, samples were analyzed in backscatter mode to enhance contrast between oomycete and liverwort tissue.

### Histochemical Staining

Callose staining was performed by gently rinsing whole plants in 0.07 M phosphate buffer (pH 9), then submerging whole plants in freshly prepared 0.05% aniline blue (w/v) solution (dissolved in 0.07 M phosphate buffer, pH 9). Plants were incubated in staining solution for 60 minutes and were imaged immediately. A total of six infection sites (two per plant; three independent plants) were assessed.

### RNA Isolation, cDNA synthesis, and qRT-PCR analysis

Total RNA was extracted from flash frozen *M. polymorpha* (TAK1) plants that were mock-inoculated (water) or infected with *P. palmivora* (ARI-td) zoospores at 1, 2, 3, or 4 days post inoculation (dpi) using the Concert Plant RNA Reagent (Invitrogen, Cat No. 12322-012) following the manufacturer’s instructions. Total RNA was extracted from axenically cultivated *P. palmivora* ARI-td using the Qiagen Plant RNeasy kit followed by on-column DNAse treatment (for RNAseq) or by using the Concert Plant RNA Reagent (for qRT-PCR analysis). All samples extracted using the Concert Plant RNA Reagent were treated with Turbo-DNAse reagent (Ambion) to degrade residual DNA contamination prior to further use. cDNA was synthesized with SuperScript II reverse transcriptase (Invitrogen) using 2 μg of total RNA following the manufacturer’s instructions. All cDNA samples were diluted 10x with nuclease free water and stored at - 20 °C until further use. qRT-PCR analyses were carried out with 2.5 μl of diluted cDNA and LightCycler 480 SYBR Green I master mix in a 10 μl volume, according to the manufacturer’s instructions. Primers for qRT-PCR analyses were designed using Primer3 [59,60] and are listed in Table S4. Specificity was validated by analyzing melt curves after each run. Three technical replicates were analysed for each of the three independent sample replicates for any given time point/treatment. Calculations of expression levels normalized to internal controls and statistical analyses were performed using R software. Graphs were generated in GraphPad Prism6.

### Cloning and Marchantia Transformation

The *Marchantia MpRabG3c* and *MpRabA1e* homologs were first identified by BlastP analysis using homologous sequences from *N. benthamiana*. Full length *MpRabG3c* (Mapoly0946s0001) and *MpRabA1e* (Mapoly0018s0008) were cloned from cDNA generated from untreated, 3 week-old *M. polymorpha* (TAK1) plants using Phusion DNA polymerase (NEB) and gateway compatible primers described in Table S4. Full length attB sequences were generated using universal attB primers that allow for N-terminal fusions. Amplicons were recombined into pDONR221 using BP clonase II (Invitrogen) following manufacturer’s instructions. Sequence verified entry clones were then recombined into pMpGWB105 [61] using LR clonase II enzyme mix (Invitrogen) following manufacturer’s instructions. Completed destination constructs were transformed into *Agrobacterium tumefaciens* GV3101 (pMP90) by electroporation. *M. polymorpha* transformation was carried out using the *Agrobacterium*-mediated thallus regeneration method using TAK1 plants [62]. Transformants were selected on solidified ½ MS media (pH 5.6) supplemented with hygromycin B (15-25 μg mL^−1^) and cefotaxime (125 μg mL^−1^). Stable, non-chimeric transgenic plants were obtained by propagating gemmae from T1 thalli.

### Library preparation and sequencing

mRNAs from *M. polymorpha* plants infected with *P. palmivora* at 3 dpi and 4 dpi, and *P. palmivora* mRNAs from the mycelium sample were purified using Poly(A) selection from total RNA sample, and then fragmented. cDNA library preparation was performed with the TruSeq^®^ RNA Sample Preparation Kit (Illumina, US) according to the manufacturer’s protocol. cDNA sequencing of the 9 samples (3 dpi, 4 dpi and mycelium sample, all in triplicates) was performed with Illumina NextSeq 2500 in 100 paired end mode. Samples were de-multiplexed and analyzed further. The raw fastq data are accessible at http://www.ncbi.nlm.nih.gov/sra/ with accession number SRP115544.

### Expression analysis

Raw reads after quality control with FastQC (https://www.bioinformatics.babraham.ac.uk/projects/fastqc/) were aligned back to the *P. palmivora* reference genome [63] using STAR (version 2.5.2b) aligner. Raw counts were obtained with FeatureCounts [64], only uniquely mapped and properly paired reads were considered further. Differentially expressed genes were identified with DESeq2 Bioconductor package [65] following pair-wise comparisons between *in planta* and mycelium samples. Differentially expressed genes (absolute LFC ≥ 2 and adjusted *p*-value ≤ 10^−3^) were used to perform hierarchical clustering of samples. Heatmaps for the differentially expressed genes were generated using R pheatmap package [66]. For the final heatmaps rlog-transformed counts median-centered by gene were used. Scripts used to analyse *P. palmivora* RNA-seq dataset and visualise differentially expressed genes are available in https://github.com/gogleva/public-palmivora.

### Secretome prediction

Putative secreted *P. palmivora* proteins were predicted and manually curated as described previously [28] based on the gene models from [63].

## ABBREVIATIONS

DEG: differentially expressed genes
DPI: days post inoculation
LFC: log fold change
FDR: false discovery rate
ORF: open reading frame
NPP: necrosis inducing *Phytophthora* protein
SCR: small cysteine-rich peptides
CWDE: cell wall degrading enzyme
PI: protease inhibitor
EPI: extracellular protease inhibitor
TM: transmembrane domain
SP: signal peptide
SYP: syntaxin

## ACKNOWLEDGMENTS

For providing materials, we thank Dr. Takashi Ueda (University of Tokyo, Japan), Dr. Jim Haseloff (University of Cambridge, UK), and Dr. Pierre-Marc Delaux (Paul Sabatier University, France). For access to *Marchantia polymorpha* genomic information (https://phytozome.jgi.doe.gov), we thank our colleagues at the Joint Genome Institute (JGI); the work conducted by the U.S. Department of Energy (DOE) JGI, a DOE Office of Science User Facility, is supported by the Office of Science of the U.S. DOE under contract DE-AC02-05CH11231. For assistance with cryo-SEM analysis, we thank Dr. Raymond Wightman (Sainsbury Laboratory, University of Cambridge). We thank Dr. Sophien Kamoun (The Sainsbury Laboratory, Norwich) for critically assessing an early draft of this manuscript. For additional critical and technical support, we thank Dr. Edouard Evangelisti, Dr. Ruth Le Fevre, Dr. Temur Yunusov, Lara Busby, and all members of the Schornack group.

## SUPPORTING INFORMATION

**Fig S1. Rhizoid inoculations do not reliably lead to colonization**

**(A)** Disease symptoms at 7 days post inoculation (dpi) of 3-week-old *M. polymorpha* (TAK1) plants that were inoculated with *P. palmivora* ARI-td zoospores or water directly onto rhizoids.

**(B)** Confocal fluorescence microscopy demonstrating *P. palmivora* ARI-td growth on 3-week-old *M. polymorpha* rhizoids from 1-3 dpi. Micrographs represent z-stack projections of merged bright field and tdTomato channels. Scale bars = 50 μm.

**Fig S2. *P. palmivora* colonizes TAK1 thalli.** Confocal fluorescence microscopy demonstrating *P. palmivora* ARI-td growth on 3-week-old *M. polymorpha* thalli from 1-4 days post inoculation (dpi). Micrographs represent z-stack projections. The merged micrographs display red pathogen fluorescence (tdTomato) overlaid on top of chlorophyll autofluorescence (turquoise). Scale bars = 100 μm.

**Fig S3. *P. palmivora* hyphae access the storage region during necrotrophy.**

Confocal fluorescence microscopy demonstrating the co-occurrence of *P. palmivora* ARI-td hyphae and necrotrophic disease symptoms in the storage region of *M. polymorpha* thalli at 7 days post inoculation (dpi). Red pathogen fluorescence is merged with bright field images. Micrographs represent z-stacked images. Scale bars = 100 μm.

**Fig S4. Biotrophic colonization of the photosynthetic layer in *Marchantia paleacea* and *Lunularia* cruciata.**

**(A)** Confocal fluorescence microscopy of sectioned thalli of *M. paleacea* and *L cruciata* colonized by *P. palmivora* ARI-td at 7 days post inoculation (dpi). Z-stacked images represent red pathogen fluorescence (tdTomato) merged with the bright field channel. Colonized air chambers are denoted by arrows. Scale bars = 100 μm

**(B)** Confocal fluorescence microscopy demonstrating haustoria morphology in *P. palmivora* ARI-td-colonized *M. paleacea* (*Mpal*) and *L. cruciata* (*Lc*) thalli at 3 dpi. Z-stacked images display red pathogen fluorescence (ARI-td) merged with plastid autofluorescence. Scale bars = 10 μm.

**Fig S5. *P. palmivora* strains vary in aggressiveness in TAK1**.

Disease symptoms of *M. polymorpha* TAK1 plants inoculated with water (mock) or zoospores of *P. palmivora* strains 7 days post inoculation (dpi). Images displayed are representative of consistent symptoms observed from 8-16 infected plants per strain.

**Fig S6. *P. infestans* does not cause disease symptoms on TAK1**

Disease progression of *M. polymorpha* TAK1 plants inoculated with water, *P. palmivora* (ARI-td) zoospores or *P. infestans* (*Pi88069*-td) zoospores. Images display consistent disease symptoms (n=8) at 1, 3, 5, and 7 days post inoculation (dpi).

**Fig S7. Validation of colonization-induced *P. palmivora* genes**

qRT-PCR analysis of *P. palmivora* (ARI-td) genes identified by RNA-seq analyses. Expression levels were quantified in an axenically propagated MZ (mycelia + zoospores) sample and during the colonization of *M. polymorpha* thalli from 1-4 days post inoculation (dpi). Expression levels were quantified relative to internal controls *PpEF1a* and *PpWS21*. Different letters indicate statistically significant differences in expression levels (ANOVA, Tukey’ HSD, p < 0.05). Performed twice with similar results.

**Fig S8. Callose does not envelope *P. palmivora* infection structures**

Confocal fluorescence microscopy of aniline blue stained *M. polymorpha* TAK1 thalli infected with *P. palmivora* ARI-td at 3 days post inoculation (dpi). Images represent z-stack projections displaying red fluorescence from the pathogen (tdTomato), callose deposition through aniline blue staining (white), bright field, or tdTomato merged with aniline blue. Asterisks (*) denote intracellular infection structures that are not enveloped by callose.

**Fig S9. Subcellular localization of MpRabA1e and MpRabG3c**

**(A)** Confocal fluorescence microscopy showing subcellular localization patterns of MpRabA1e in 35S:mCitrine-MpRabA1e/TAK1 gemmae. Micrographs display mCitrine fluorescence, plastid autofluorescence (magenta), both channels merged, or bright field images. Scale bars = 10 μm.

**(B)** Confocal fluorescence microscopy showing subcellular localization patterns of MpRabG3c in 35S:mCitrine-MpRabG3c/TAK1 gemmae. Micrographs display mCitrine fluorescence, plastid autofluorescence (magenta), both channels merged, or bright field images. Scale bars = 10 μm.

**S1 Table. Colonization-induced *P. palmivora* genes** (abs(LFC) >= 2, p-value≤10^−3^)

**S2 Table. Plants used in this study**.

**S3 Table. Pathogen strains used in this study**

**S4 Table. Primers used in this study**.

## REFERENCES

[1] Remy W, Taylor TN, Hass H, Kerp H. Four hundred-million-year-old vesicular arbuscular mycorrhizae. Proc. Natl. Acad. Sci. USA 1994; 91(25): 11841–11843.

[2] Taylor TN, Remy W, Hass H, Kerp H. Fossil arbuscular mycorrhizae from the early devonian. Mycologia 1995; 87(4): 560–73.

[3] Krings M, Taylor TN, Hass H, Kerp H, Dotzler N, Hermsen EJ. Fungal endophytes in a 400-million-yr-old land plant: infection pathways, spatial distribution, and host responses. New Phytologist 2007; 174(3): 648–57.

[4] Strullu-Derrien C, Kenrick P, Pressel S, Duckett JG, Rioult J-P, Strullu D-G. Fungal associations in Horneophyton ligneri from the Rhynie Chert (c. 407 million year old) closely resemble those in extant lower land plants: novel insights into ancestral plant-fungus symbioses. New Phytologist 2014; 203(3): 964–79.

[5] Ligrone R, Carafa A, Lumini E, Bianciotto V, Bonfante P, Duckett JG. Glomeromycotean associations in liverworts: a molecular, cellular, and taxonomic analysis. American Journal of Botany 2007; 94(11): 1756–77.

[6] Adams DG, Duggan PS. Cyanobacteria-bryophyte symbioses. Journal of Experimental Botany 2008; 59(5): 1047–58.

[7] Pressel S, Bidartondo MI, Ligrone R, Duckett JG. Fungal symbioses in bryophytes: new insights in the twenty first century. Phytotaxa 2010; 9(30): 238–53. doi: http://dx.doi.org/10.11646/phytotaxa.9.1.13

[8] Desiro A, Duckett JG, Pressel S, Villarreal JC, Bidartondo MI. Fungal symbioses in hornworts: a chequered history. Proc Biol Sci 2013; 280(1759): 20130207. doi: http://dx.doi.org/10.1098/rspb.2013.0207

[9] Wang B, Yeun LH, Xue J-Y, Liu Y, Ane J-M, Qiu Y-L. Presence of three mycorrhizal genes in the common ancestor of land plants suggests a key role of mycorrhizas in the colonization of land by plants. New Phytologist 2010; 186(2): 514–25.

[10] Delaux P-M, Radhakrishnan GV, Jayaraman D, Cheema J, Malbreil M, Volkening JD, Sekimoto H, Nishiyama T, Melkonian M, Pokorny L, Rothfels CJ, Sederoff HW, Stevenson DW, Surek B, Zhang Y, Sussman MR, Dunand C, Morris RJ, Roux C, Wong G, Oldroyd G, Ane J-M. Algal ancestor of land plants was preadapted for symbiosis. Proc. Natl. Acad. Sci. USA 2015; 112(43): 13390–13395.

[11] Petre B, Kamoun S. How do filamentous pathogens deliver effector proteins into plant cells? PLoS Biology 2014; 12(2):e1001801. doi: 10.1371/journal.pbio.1001801.

[12] Roth R, Paszkowski U. Plant carbon nourishment of arbuscular mycorrhizal fungi. Current Opinion in Plant Biology 2017; 39: 50–56

[13] Parniske M. Intracellular accommodation of microbes by plants: a common developmental program for symbiosis and disease? Current Opinion in Plant Biology 2000; 3(4): 320–28.

[14] Rey T, Schornack S. Interactions of beneficial and detrimental root-colonizing filamentous microbes with plant hosts. Genome Biology 2013; 14(6): 121. doi: 10.1186/gb-2013-14-6-121.

[15] Ivanov S, Fedorova E, Bisseling T. Intracellular plant microbe associations: secretory pathways and the formation of perimicrobial compartments. Current Opinion in Plant Biology 2010; 13(4): 372–77

[16] Zeilinger S, Gupta VK, Dahms TES, Silva RN, Singh HB, Upadhyay RS, Gomes EV, Tsui CKM, Nayak SC. Friends or foes? Emerging insights from fungal interactions with plants. FEMS Microbiology Reviews 2016; 40(2): 182–207

[17] Qiu YL, Li L, Wang B, Chen Z, Knoop V, Groth-Molanek M, Dombrovska O, Lee J, Kent L, Rest L, Estabrook GF, Hendry TA, Taylor DW, Testa CM, Ambros M, Crandall-Stotler B, Duff RJ, Stech M, Frey W, Quandt D, Davis CC. The deepest divergences in land plants inferred from phylogenomic evidence. Proceedings of the National Academy of Sciences USA 2006; 103(42): 15511–6

[18] Renzaglia KS, Schuette S, Duff RJ, Ligrone R, Shaw AJ, Mishler BD, Duckett JG. Bryophyte phylogeny: advancing the molecular and morphological frontiers. The Bryologist 2007; 110(2): 179–213.

[19] Cox CJ, Li B, Foster PG, Embley TM, Civan P. Conflicting phylogenies for early land plants are caused by composition biases among synonymous substitutions. Systematic Biology 2014; 63(2): 272–79

[20] Ruhfel BR, Gitzendanner MA, Soltis PS, Soltis DE, Burleigh JG. From algae to angiosperms-inferring the phylogeny of green plants (Viridiplantae) from 360 plastid genomes. BMC Evolutionary Biology 2014; 14: 23. doi: 10.1186/1471-2148-14-23.

[21] Ponce de Leon I. The moss Physcomitrella patens as a model system to study interactions between plants and phytopathogenic fungi and oomycetes. J. Pathogens 2011; 2011:719873. doi: 10.4061/2011/719873

[22] Overdijk EJ, De Keijzer J, De Groot D, Schoina C, Bouwmeester K, Ketelaar T, Govers F. Interaction between the moss Physcomitrella patens and Phytophthora: a novel pathosystem for live-cell imaging of subcellular defence. J. Microscopy 2016; 263(2): 171–80

[23] Ponce de Leon I, Montesano M. Adaptation mechanisms in the evolution of moss defenses to microbes. Frontiers in Plant Science 2017; 8: 366. doi: 10.3389/fpls.2017.00366

[24] Ishizaki K, Nishihama R, Yamato KT, Kohchi T. Molecular genetic tools and techniques for Marchantia polymorpha research. Plant and Cell Physiology 2016; 57(2): 262–70

[25] Shimamura M. Marchantia polymorpha: taxonomy, phylogeny, and morphology of a model system. Plant and Cell Physiology 2016; 57(2): 230–56.

[26] Rey T, Chatterjee A, Buttay M, Toulotte J, Schornack S. Medicago truncatula symbiosis mutants affected in the interaction with a biotrophic root pathogen. New Phytologist 2015; 206(2): 385–95.

[27] Le Fevre R, O’Boyle B, Moscou MJ, Schornack S. (2016) Colonization of barley by the broad-host hemibiotrophic pathogen Phytophthora palmivora uncovers a leaf development-dependent involvement of Mlo. Molecular Plant Microbe Interactions 2016; 29(5): 385–95.

[28] Evangelisti E, Gogleva A, Hainaux T, Doumane M, Tulin F, Quan C, Yunusov T, Floch K, Schornack S. Time-resolved dual root-microbe transcriptomics reveals early induced Nicotiana benthamiana genes and conserved infection-promoting Phytophthora palmivora effectors. BMC Biology 2017; 15(1): 39. doi: 10.1186/s12915-017-0379-1

[29] Judelson HS, Blanco FA. The spores of Phytophthora: weapons of the plant destroyer. Nat. Rev. Microbiol. 2005; 3(1): 47–58.

[30] Morgan W, Kamoun S. RXLR effectors of plant pathogenic oomycetes. Current Opinion in Microbiology 2007; 10(4): 332–38.

[31] Anderson RG, Deb D, Fedkenheuer K, McDowell JM. Recent progress in RXLR effector research. Molecular Plant-Microbe Interactions 2015; 28(10): 1063–72.

[32] Wang S, Boevink PC, Welsh L, Zhang R, Whisson SC, Birch PRJ. Delivery of cytoplasmic and apoplastic effectors from Phytophthora infestans haustoria by distinct secretion pathways. New Phytologist 2017; 216(1): 205–215

[33] Hardham AR. Cell biology of plant-oomycete interactions. Cellular Microbiology 2007; 9(1): 31–9

[34] Torres GA, Sarria GA, Martinez G, Varon F, Drenth A, Guest D. Bud rot caused by Phytophthora palmivora: a destructive emerging disease of oil palm. Phytopathology 2016; 106(4): 320–9.

[35] Avrova AO, Boevink PC, Young V, Grenville-Briggs LJ, van West P, Birch PR, Whisson SC. A novel Phytophthora infestans haustorium-specific membrane protein is required for infection of potato. Cellular Microbiology 2008; 10(11): 2271–84

[36] Ah Fong AMV, Judelson HS. Cell cycle regulator Cdc14 is expressed during sporulation but not hyphal growth in the fungus-like oomycete Phytophthora infestans. Molecular Microbiology 2003; 50(2): 487–94.

[37] Bellincampi D, Cervone F, Lionetti V. Plant cell wall dynamics and wall-related susceptibility in plant-pathogen interactions. Frontiers in Plant Science 2014; 5: 228 doi: 10.3389/fpls.2014.00226

[38] Faulkner C. A cellular backline: specialization of host membranes defence. Journal of Experimental Botany 2015; 66(6): 1565–71.

[39] Bozkurt TO, Belhaj K, Dagdas YF, Chaparro-Garcia A, Wu CH, Cano LM, Kamoun S. Rerouting of plant late endocytic trafficking toward a pathogen interface. Traffic 2015; 16(2): 204–26.

[40] Lu YJ, Schornack S, Spallek T, Geldner N, Chory J, Schellmann S, Schumacher K, Kamoun S, Robatzek S. Patterns of plant subcellular responses to successful oomycete infections reveal differences in host cell reprogramming and endocytic trafficking. Cell Microbiology 2012; 14(5): 682–97.

[41] Zhang C, Hicks GR, Raikhel NV. Plant vacuole morphology and vacuolar trafficking. Frontiers in Plant Science 2014; 5: 476. doi: 10.3389/fpls.2014.00476

[42] Gu Y, Zavaliev R, Dong X. Membrane trafficking in plant immunity. Molecular Plant 2017; 10(8): 1026–1034

[43] Kanazawa T, Ueda T. Exocytic trafficking pathways in plants: why and how they are redirected. New Phytologist 2017; 215(3): 952–57

[44] Kanazawa T, Era A, Minamino N, Shikano Y, Fujimoto M, Uemura T, Nishihama R, Yamato KT, Ishizaki K, Nishiyama T, Kohchi T, Nakano A, Ueda T. SNARE molecules in Marchantia polymorpha: unique and conserved features of the membrane fusion machinery. Plant and Cell Physiology 2016; 57(2): 307–24.

[45] Ishizaki K, Mizutani M, Shimamura M, Nishihama R, Kohchi T. Essential role of the E3 ubiquitin ligase nopperabo1 in schizogenous intercellular space formation in the liverwort Marchantia polymorpha. Plant Cell 2013; 25(10): 4075–84

[46] Schussler A. Glomus claroideum forms an arbuscular mycorrhiza-like symbiosis with the hornwort Anthoceros punctatus. Mycorrhiza 2000; 10(1): 15–21

[47] McLachlan DH, Kopischke M, Robatzek S. Gate control: guard cell regulation by microbial stress. New Phytologist 2014; 203(4): 1049–63.

[48] Melotto M, Zhang L, Oblessuc PR, He S-Y. Stomatal defense a decade later. Plant Physiology 2017; 174(2): 561–71.

[49] Field KJ, Duckett JG, Cameron DD, Pressel S. Stomatal density and aperture in non-vascular land plants are non-responsive to above-ambient atmospheric CO2 concentrations. Annals of Botany 2015; 115(6): 915–22

[50] Sarkar P, Bosneaga E, Auer M. Plant cell walls throughout evolution: towards a molecular understanding of their design principles. Journal of Experimental Botany 2009; 60(13): 3615–35

[51] Takikawa Y, Senga Y, Nonomura T, Matsuda Y, Kakutani K, Toyoda H. Targeted destruction of fungal structures of Erysiphe trifoliorum on flat leaf surfaces of Marchantia polymorpha. Plant Biology 2014; 16(1): 291–5.

[52] Latijnhouwers M, de Wit PJGM, Govers F. Oomycetes and fungi: similar weaponry to attack plants. Trends in Microbiology 2003; 11(10): 462–69.

[53] Wang E, Schornack S, Marsh JF, Gobbato E, Schwessinger B, Eastmond P, Schultze M, Kamoun S, Oldroyd GED. A common signaling process that promotes mycorrhizal and oomycete colonization of plants. Current Biology 2012; 22(23): 2242–46.

[54] Ligrone R, Duckett JG, Renzaglia KS. Major transitions in the evolution of early land plants: a bryological perspective. Annals of Botany 2012; 109(5): 851–71

[55] Catalano CM, Czymmek KJ, Gann JG, Sherrier DJ. Medicago truncatula syntaxin SYP132 defines the symbiosome membrane and infection droplet membrane in root nodules. Planta 2007; 225(3): 541–50

[56] Pan H, Oztas O, Zhang X, Wu X, Stonoha C, Wang E, Wang B, Wang D. A symbiotic SNARE protein generated by alternative termination of transcription. Nature Plants 2016; 2: 15197. doi: 10.1038/nplants.2015.197

[57] Caillaud M-C, Wirthmueller L, Sklenar J, Findlay K, Piquerez SJM, Jones AME, Robatzek S, Jones JDG, Faulkner C. The plasmodesmatal protein PDLP1 localises to haustoria-associated membranes during downy mildew infection and regulates callose deposition. PLOS Pathogens 2014; 10(10): e1004496. doi: https://doi.org/10.1371/journal.ppat.1004496

[58] Wightman R, Wallis S, Aston P. Hydathode pit development in the alpine plant Saxifraga cochlearis. Flora 2017; 233: 99–108. doi: https://doi.org/10.1016/j.flora.2017.05.018

[59] Koressaar T, Remm M. Enhancements and modifications of primer design program Primer3. Bioinformatics 2007; 23(10): 1289–91

[60] Untergasser A, Cutcutache I, Koressaar T, Ye J, Faircloth BC, Remm M, Rozen SG. Primer3 – new capabilities and interfaces. Nucleic Acids Research 2012; 40(15): e115. doi: 10.1093/nar/gks596

[61] Ishizaki K, Nishihama R, Ueda M, Inoue K, Ishida S, Nishimura Y, Shikanai T, Kohchi T. Development of gateway binary vector series with four different selection markers for the liverwort Marchantia polymorpha. PLoS ONE 2015; 10(9): e0138876. doi: https://doi.org/10.1371/journal.pone.0138876

[62] Kubota A, Ishizaki K, Hosaka M, Kohchi T. Efficient agrobacterium-mediated transformation of the liverwort Marchantia polymorpha using regenerating thalli. Biosci. Biotechnol. Biochem. 2013; 77(1): 167–77

[63] Ali SS, Shao J, Lary DJ, Kronmiller BA, Shen D, Strem MD, Amoako-Attah I, Akrofi AY, Begoude BAD, ten Hoopen GM, Coulibaly K, Kebe BI, Melnick RL, Guiltinan MJ, Tyler BM, Meinhardt LW, Bailey BA. Phytophthora megakarya and Phytophthora palmivora, closely related causal agents of cacao black pod rot, underwent increases in genome sizes and gene numbers by different mechanisms. Genome Biology and Evolution 2017; 9(3): 536–57

[64] Liao Y, Smyth GK, Shi W. featureCounts: an efficient general purpose program for assigning sequence reads to genomic features. Bioinformatics 2014; 30(7): 923–30

[65] Love MI, Huber W, Anders S. Moderated estimation of fold change and dispersion for RNA-seq data with DESeq2. Genome Biology 2014; 15(12): 550 https://doi.org/10.1186/s13059-014-0550-8

[66] Kolde R. pheatmap: Pretty Heatmaps. R package version 1.0.8. 2015

